# Presynaptic control of top-down signaling in neocortical layer 1

**DOI:** 10.64898/2026.01.31.703032

**Authors:** Kelly E. Bonekamp, M. Lea Ratz, Grant R. Gillie, Lingxi Xiong, Mya K. Sebek, Shane R. Crandall

## Abstract

The integration of bottom-up and top-down signals is crucial for perception and behavior. Neocortical layer 1 (L1) is a key target for top-down inputs and contains various GABAergic interneurons. Here, we investigated whether GABA can presynaptically regulate neurotransmitter release at top-down synapses targeting L1 of primary somatosensory cortex (S1). Our findings show that presynaptic GABA_B_ receptor activation suppresses corticocortical inputs from primary motor (M1) and secondary somatosensory cortices (S2) more than those from the posterior medial nucleus of the thalamus (POm). This effect varied by target cell type, with GABAergic interneurons being less affected. Finally, we demonstrate that L1 neuron-derived neurotrophic factor (NDNF)-expressing interneurons inhibit top-down synapses more effectively than somatostatin-expressing interneurons, and that POm drives L1 neurogliaform cells, a subtype of NDNF interneuron that elicits unitary GABA_B_ responses. These results reveal a novel circuit in which higher-order thalamus influences top-down corticocortical communication via L1 neurogliaform cells.

## INTRODUCTION

One of the remarkable features of the brain is its ability to perceive and adapt behavior according to immediate demands and changing environments. This context-dependent process relies on the continuous integration of bottom-up external sensory cues and top-down internal signals that reflect ongoing expectations, predictions, motor actions, and motivations^1–9^. Neural circuits in the neocortex play a crucial role in this process, contributing significantly to normal sensory perception, learning, and consciousness^10–15^.

Layer 1 (L1) of the neocortex, long considered mysterious, is now thought to be critical for integrating bottom-up and top-down information^13,16–19^. This unique layer lacks excitatory cells and comprises only diverse GABAergic inhibitory cells and the distal tuft dendrites of pyramidal neurons^20,21^. L1 is also a major target of top-down afferents originating from several other brain structures, including other cortical areas^22–25^ and higher-order thalamic regions^26,27^, as well as subcortical neuromodulatory centers^28–33^. Together, these afferent projections convey diverse contextual signals to L1^34–36^, which are received and processed by the distal tuft dendrites before being integrated with more proximal bottom-up inputs, thereby changing the gain of pyramidal neurons^37–39^. The unique structural organization of L1 suggests that its inhibitory cells may play an essential role in contextual processing by controlling the communication of top-down signals to the distal tuft dendrites of pyramidal cells. However, a complete understanding of the inhibitory circuit mechanisms that regulate this integration is currently lacking.

While the role of distinct GABAergic interneurons in regulating signal integration in pyramidal cells through dendritic inhibition and disinhibition is well established^39–48^, GABA can also act presynaptically to control neurotransmitter release^49–53^. Presynaptic inhibition typically depends on GABA_B_ receptor activation^49–51^, and is widely used throughout the nervous system to serve various functions, including stabilizing neural networks, adjusting synaptic gain, relieving synaptic depression, and promoting synaptic plasticity^54–56^. Given the diversity of inhibitory interneurons in the neocortex and the various ways they shape cortical activity^57,58^, it is important to determine whether presynaptic GABA_B_ receptors suppress top-down signaling and which cell type may be responsible. Understanding this is critical for fully comprehending how the integration of bottom-up and top-down signals is modulated by inhibition. Remarkably, compelling support for this concept comes from studies showing that distinct types of cortical interneurons mediate GABA_B_ inhibition^59–61^.

Interneurons expressing neuron-derived neurotrophic factor (NDNF) form a major component of L1, and one of the subtypes of NDNF interneurons includes neurogliaform (NGF) cells^62^. NGF cells feature a very dense local axon arborization with an unusually high number of synaptic boutons that are positioned further from their targets than usual, resulting in GABA volume transmission^60,63^. They are also the only class of inhibitory interneurons that activate GABA_B_ receptors at both postsynaptic and presynaptic sites with a single action potential^59,60,63^. Collectively, the presence of NGF cells and the density of top-down synaptic inputs in L1 suggest that NGF cells could be key to controlling top-down signaling via activation of presynaptic GABA_B_ receptors. Indeed, Pardi et al. (2020) showed that output from NDNF cells controls afferent signaling from the higher-order auditory thalamus to the secondary auditory cortex via presynaptic GABA_B_ receptors^61^.

Here, we combined *in vitro* electrophysiology and optogenetic strategies to ask whether other glutamatergic afferents to cortical L1 are subject to presynaptic modulation by GABA_B_ receptors and to identify the underlying circuit elements that may mediate this inhibition in L1. Using the mouse vibrissal primary somatosensory cortex (S1), we demonstrate that the synaptic strength of L1 targeting cortical inputs originating from vibrissal primary motor (M1) and vibrissal secondary somatosensory cortices (S2) is strongly suppressed by GABA_B_ receptor activation. In contrast, inputs from the higher-order posterior medial nucleus (POm) of the thalamus are modulated considerably less. We also found that NDNF cell output is sufficient to suppress M1 synapses via presynaptic GABA_B_ receptor activation and that POm neurons selectively drive NGF cells in L1. Taken together, our results describe a new circuit in which inputs from higher-order thalamus influence top-down corticocortical communication by activating NGF cells, which, in turn, suppress L1 corticocortical transmission by activating presynaptic GABA_B_ receptors.

## RESULTS

### GABA_B_ receptor activation suppresses top-down projections targeting L1

We initially assessed the efficacy of GABA_B_ receptor activation on glutamatergic transmission from long-range M1, S2, and POm axons in L2 pyramidal neurons of S1 in acute brain slices. All three of these pathways terminate in L1 of S1 and synapse on L2/3 cells^45,64–66^, where they are thought to modulate the sensory responsiveness of these and other pyramidal cells by conveying contextual information related to voluntary whisker control^38,67,68^, behavioral state^36,69–71^, and stimulus orientation^72^. To selectively activate these pathways using an optogenetic approach, we injected M1, S2, or POm with an adeno-associated virus (AAV2) carrying a transgene encoding the light-gated cation channel Channelrhodopsin-2 fused to enhanced yellow fluorescent protein (ChR2/EYFP)^73–75^. These injections resulted in robust ChR2/EYFP expression in M1, S2, and POm cells, as well as in their terminal arbors in S1, where they densely innervated L1, consistent with previous reports^76–80^ (**Fig. 1a, 1c, and 1e**).

**Fig 1.**
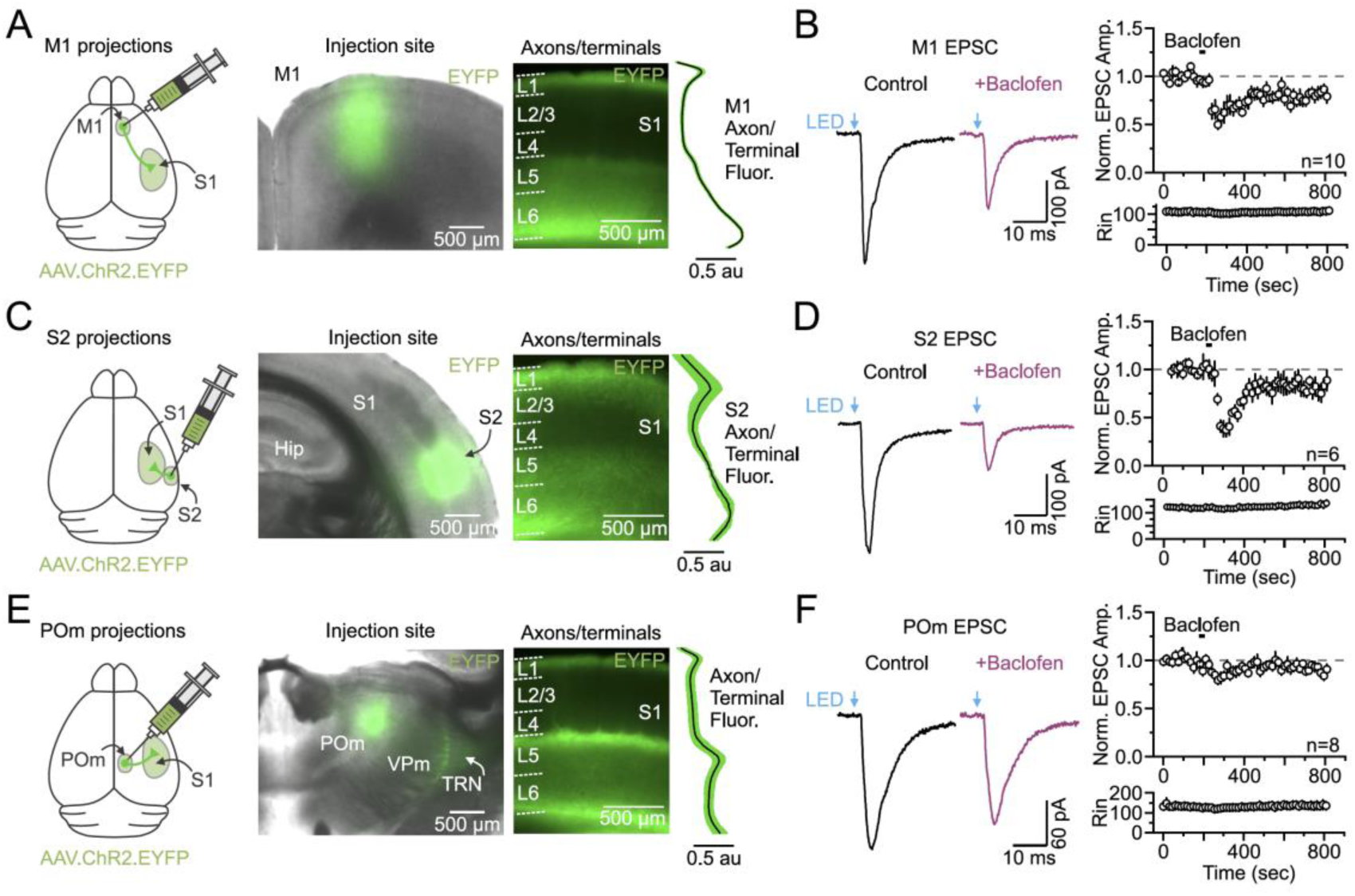
GABA_B_ receptor activation inhibits the synaptic strength of L1 top-down corticocortical inputs more than thalamic inputs. **(a)** Left, schematic showing a viral injection into the M1 of an AAV carrying transgenes for ChR2/EYFP. Middle, a merged brightfield image of a live slice and EYFP fluorescence showing the location of the viral injection into M1. Right, fluorescence image of a different live slice from the same brain showing EYFP-expressing M1 axons/terminals in S1, and the mean normalized fluorescent intensity profile as a function of depth from pia (n = 13 slices, 13 mice). (**b)** Left, average M1 synaptic response in a L2 pyramidal cell of S1 evoked by a brief (0.5 ms), single optical stimulus in control conditions (black) and after adding baclofen (2.5 µM, magenta). Right, population data showing the average time course of baclofen effects (30 sec application) on M1 evoked EPSC amplitudes and postsynaptic input resistance (n = 10 cells, 8 mice). Data are normalized to the 3 min mean baseline amplitude before baclofen application. Only the cells with stable recordings throughout the entire 15 min are plotted. **(c)** Same as **(a)** but of AAV.ChR2.EYFP injections into S2 (n = 3 slices, 3 mice). **(d)** Same as **(b)** but of baclofen effects on S2 synaptic responses (n = 6 cells, 3 mice). **(e),** Same as (**a)** but of AAV.ChR2.EYFP injections into POm (n = 6 slices, 6 mice). **(f)** Same as **(b)** but of baclofen effects on POm synaptic responses (n = 8 cells, 3 mice). Hip, hippocampus, TRN, thalamic reticular nucleus, VPm, ventral posterior medial nucleus.

Brief optical stimulation (0.5 ms) of M1, S2, or POm axons/terminals evoked short-latency (∼2 ms) excitatory postsynaptic currents (EPSCs) in L2 pyramidal cells of S1 (**Fig. 1b, 1d, and 1f**). To minimize postsynaptic effects of GABA_B_ receptor activation, potassium (K^+^) channels were blocked using a cesium (Cs^+^)-based intracellular solution under voltage-clamp while holding at -75 mV. For the M1 pathway, application of the GABA_B_ receptor agonist baclofen (2.5 µM; 30 sec) significantly reduced the strength of EPSCs by 36.02 ± 6.8% (n = 13 cells, 9 mice; p = 0.0012, Wilcoxon paired signed-rank test; **Fig. 1b**). This suppression was prevented with bath application of the GABA_B_ receptor antagonist CGP-55845 (4 µM) (Baclofen only: 39.4 ± 5.4%, +CGP: 6.95 ± 5.1%; n = 4 cells, 3 mice; p = 0.0304, paired sample t-test; **Fig. S1a**). The effect was also dose dependent, with 0.31, 0.63 1.25, 2.5, 5, 10, and 20 µM baclofen reducing M1 EPSC strength to 10.3 ± 4.8%, 14 ± 5.2%, 30.1 ± 2.4%, 36 ± 6.8%, 33.8 ± 5.3%, 65.6 ± 4.6%, and 61.2 ± 5.5%, respectively (**Fig. S1b**).

When optically stimulating the S2 pathway, we found that the inhibitory effect of baclofen (2.5 µM) on the strength of S2 synapses was also robust, significantly suppressing the amplitude of EPSCs by 50.1 ± 3.8% (n = 8, 3 mice; p = 3.24 x 10^-6^, paired sample t-test; **Fig. 1d**). In contrast, the same concentration of baclofen reduced the strength of POm EPSCs by only 17.7 ± 3.2% (n = 12 cells, 3 mice; p = 4.88 x 10^-4^, Wilcoxon signed-rank pair test; **Fig. 1f**), significantly less than that observed for corticocortical synapses (M1 vs POm: p = 0.032; S2 vs POm: p = 9.66 x 10^-4^; Kruskal-Wallis ANOVA with Dunn’s post-hoc test). Application of a higher concentration of baclofen (10 µM) further suppressed POm EPSCs, but the magnitude of modulation was still less than that observed for M1 synapses at the same concentration (p = 0.00875, two-sample t-test; **Fig. S1b**).

The lack of a change in postsynaptic input resistance upon baclofen application suggests that the effects on synaptic strength are due to presynaptic GABA_B_ receptor activation (**Fig. 1b, 1d, and 1f**). Consistent with this observation, application of baclofen also significantly increased the paired-pulse facilitation ratio (PPR) and coefficient of variation (CV) of optically evoked M1 and S2 EPSCs, which are dependent on initial release probability (M1 PPR: 1.34 ± 0.06, +Baclofen: 1.54 ± 0.08, p = 8.53 x 10^-4^, paired sample t-test; M1 CV: 0.15 ± 0.01; +Baclofen: 0.23 ± 0.02; n = 13 cells, 9 mice; p = 4.22 x 10^-4^, paired sample t-test; S2 PPR: 1.17 ± 0.06; +Baclofen: 1.51 ± 0.10; p = 0.00223, paired sample t-test; S2 CV: 0.13 ± 0.2; +Baclofen: 0.31 ± 0.05; n = 8 cells, 3 mice; p = 0.00781, Wilcoxon paired signed-rank test; **Fig. 2a-2d**). Although not significant, baclofen did tend to reduce the magnitude of paired-pulse depression (i.e., increase the PPR) while also increasing the CV of optically evoked POm EPSCs, which is in line with the overall weaker modulation at these synapses (POm PPR: 0.94 ± 0.04; +Baclofen: 0.97 ± 0.04; p = 0.36641, paired sample t-test; POm CV: 0.12 ± 0.02, Baclofen CV: 0.15 ± 0.02; n = 12 cells, 3 mice; p = 0.085, 8paired sample t-test; **Fig. 2e and 2f**).

**Fig 2.**
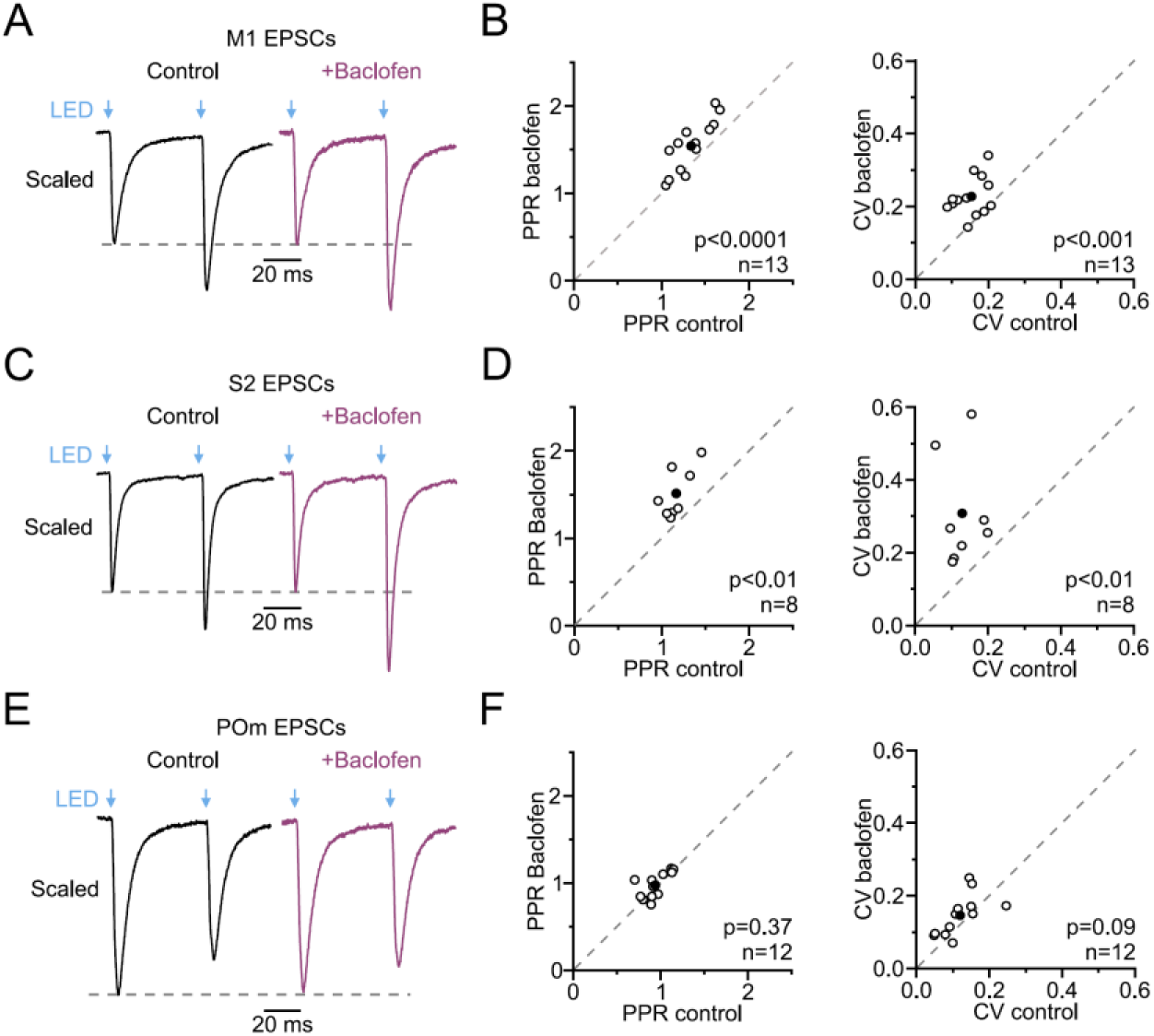
Baclofen acts presynaptically to influence the synaptic strength of top-down inputs to L1. **(a)** Left, average scaled M1 synaptic response in a L2 pyramidal cell of S1 evoked by a pair of optical stimuli delivered at 20 Hz in control conditions (black) and after adding baclofen (2.5 µM, magenta). **(b)** Population data comparing the peak PPR (left) and CV (right) in control conditions and after adding baclofen (n = 13 cells, 9 mice). The filled black circle represents the mean. **(c-d)** Same as (**a-b)** but of baclofen effects on the PPR and CV of S2 synaptic responses (n = 8 cells, 3 mice). **(e-f)** Same as (**a-b)** but of baclofen effects on the PPR and CV of POm synaptic responses (n = 12 cells, 3 mice).

Together, these results reveal that GABA_B_ receptor activation significantly decreases the strength of L1 M1, S2, and POm synapses targeting L2 pyramidal neurons in S1, and that this inhibition arises primarily from presynaptic mechanisms. Moreover, these data indicate that corticocortical and thalamocortical synapses exhibit distinct sensitivities to baclofen, with important implications for how GABA modulates top-down transmission to L1 of the neocortex.

### GABA_B_ suppression of corticocortical inputs is target cell dependent

Our results suggest that top-down corticocortical synapses are more sensitive to presynaptic GABA_B_ activation than higher-order thalamocortical synapses. In addition to L2 pyramidal cells, these corticocortical projections also synapse onto L5 pyramidal cells, which are thought to play critical roles in sensorimotor processing, decision-making, and learning^38,39,64,81–87^. To determine whether GABA_B_ receptor activation modulates these synapses, we tested the M1 pathway while recording from L5 pyramidal neurons. We found that application of baclofen (2.5 µM) had the same suppressive effect on the M1 responses of L5 as L2 cells (p = 0.58931, Mann-Whitney test), significantly reducing the synaptic strength of optically evoked EPSCs by 36.8 ± 5.5% (n = 10, 5 mice; p = 0.00195, Wilcoxon paired signed-rank test; **Fig. 3a**).

**Fig 3.**
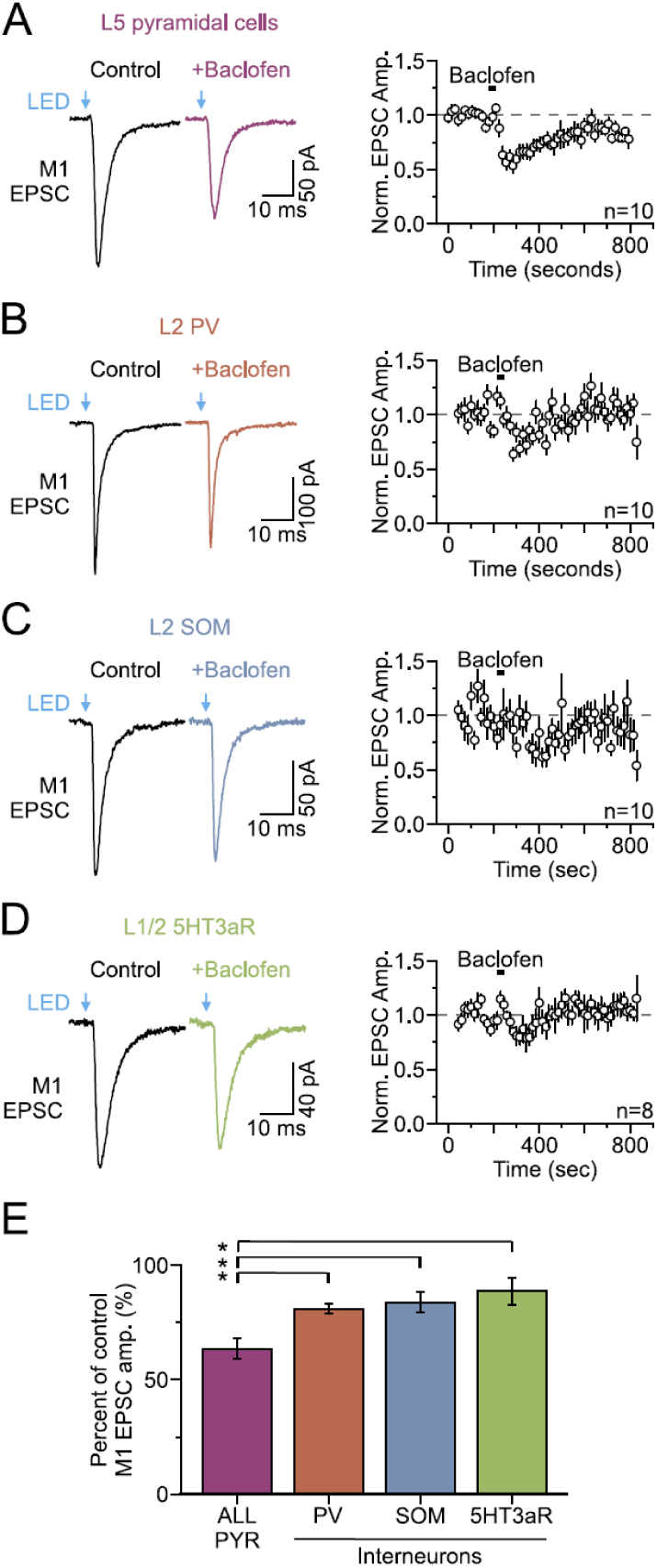
GABA_B_ receptor activation is less effective at modulating the synaptic strength of M1 input to inhibitory interneurons than to excitatory pyramidal neurons. **(a)** Left, average M1 synaptic response in a L5 pyramidal cell of S1 evoked by a single optical stimulus in control conditions (black) and after adding baclofen (2.5 µM, magenta). Right, population data showing the average time course of baclofen effects on M1-evoked EPSC amplitudes (n = 10 cells, 5 mice). **(b-d)** Left, average M1 synaptic response in a L2 PV-**(b)**, SOM-**(c)**, and 5HT3aR-**(d)** expressing inhibitory interneuron evoked by a single optical stimulus in control conditions (black) and after adding baclofen (colored traces). Right, population data showing the average time course of baclofen effects on M1-evoked EPSC amplitudes (PV: n = 10 cells, 2 mice; SOM: n = 10 cells, 6 mice; 5HT3aR: n = 8 cells, 5 mice). Data in **a-d** (right) are normalized to the 3 min mean baseline amplitude before baclofen application. **(e)** Summary plot showing the effect of baclofen (2.5 µM) as the percentage of control EPSC amplitude for excitatory pyramidal neurons (L2/5; n = 20 cells, 13 mice) and inhibitory interneurons. Asterisks denote significant differences (p < 0.05).

Corticocortical projections also target various inhibitory interneuron types that play key roles in regulating the function and dynamics of nearby excitatory cells^45,65,81^. To explore how GABA_B_ receptor activation impacts the synaptic strength of corticocortical input to inhibitory cells in superficial layers of S1, we again tested the M1 pathway using well-established transgenic mice expressing fluorescent markers for three major GABAergic cell classes in the cortex: parvalbumin (PV)-, somatostatin (SOM)-, and ionotropic serotonergic receptor (5HT3aR)-expressing interneurons^88,89^. Bath application of baclofen (2.5 µM) significantly decreased the optically evoked M1 EPSC amplitude in all three cell classes by 10-20% (PV: 19.1% ± 2.1%, n = 10, 2 mice; p = 7.93 x 10^-6^, paired sample t-test; SOM: 16.5 ± 4.1%, n = 11, 6 mice; p = 0.00236, paired sample t-test; 5HT3aR: 11.5 ± 3.6%, n = 14, 5 mice; p = 0.00703, paired sample t-test; **Fig. 3b-d**). Although there was no difference in presynaptic modulation among interneurons (p > 0.05), the magnitude of suppression was significantly less than that at excitatory pyramidal cells (p = 6.79 x 10^-4^, Kruskal-Wallis ANOVA with Dunn’s post-hoc test; PYR-PV: p = 0.049; PYR-SOM: p = 0.0089; PYR-5HT3aR: p = 0.0118; **Fig. 3e**). Together, these results strongly suggest that endogenous GABA release within S1 preferentially suppresses top-down corticocortical afferents targeting excitatory pyramidal cells.

### NDNF cells contribute to presynaptic modulation of L1 top-down synapses

What GABAergic interneuron subtype could mediate presynaptic modulation in L1 of S1? Interneurons that express NDNF are a strong candidate because their cell bodies and axons are mostly restricted to L1^62,90^, a subset of these cells are known to produce GABA_B_ responses^62,90^, and they have been shown to exert presynaptic control over L1 afferents from the higher-order auditory thalamus^61^. To test this hypothesis, we used a dual-color optogenetic approach to independently excite two distinct neural circuit elements^91,92^. Specifically, we injected a Cre-independent AAV-ChrimsonR-tdT in M1 to express the red-shifted opsin ChrimsonR in M1 axons, followed by a Cre-dependent AAV-ChR2-EYFP in the ipsilateral S1 of *NDNF-Cre* mice to express the blue-shifted opsin ChR2 in NDNF neurons (**Fig. 4a**). This strategy allowed us to monitor the strength of optically evoked M1 synaptic responses in non-expressing L2 pyramidal neurons of S1 before and after optical activation of local NDNF interneurons (**Fig. 4b**).

**Fig 4.**
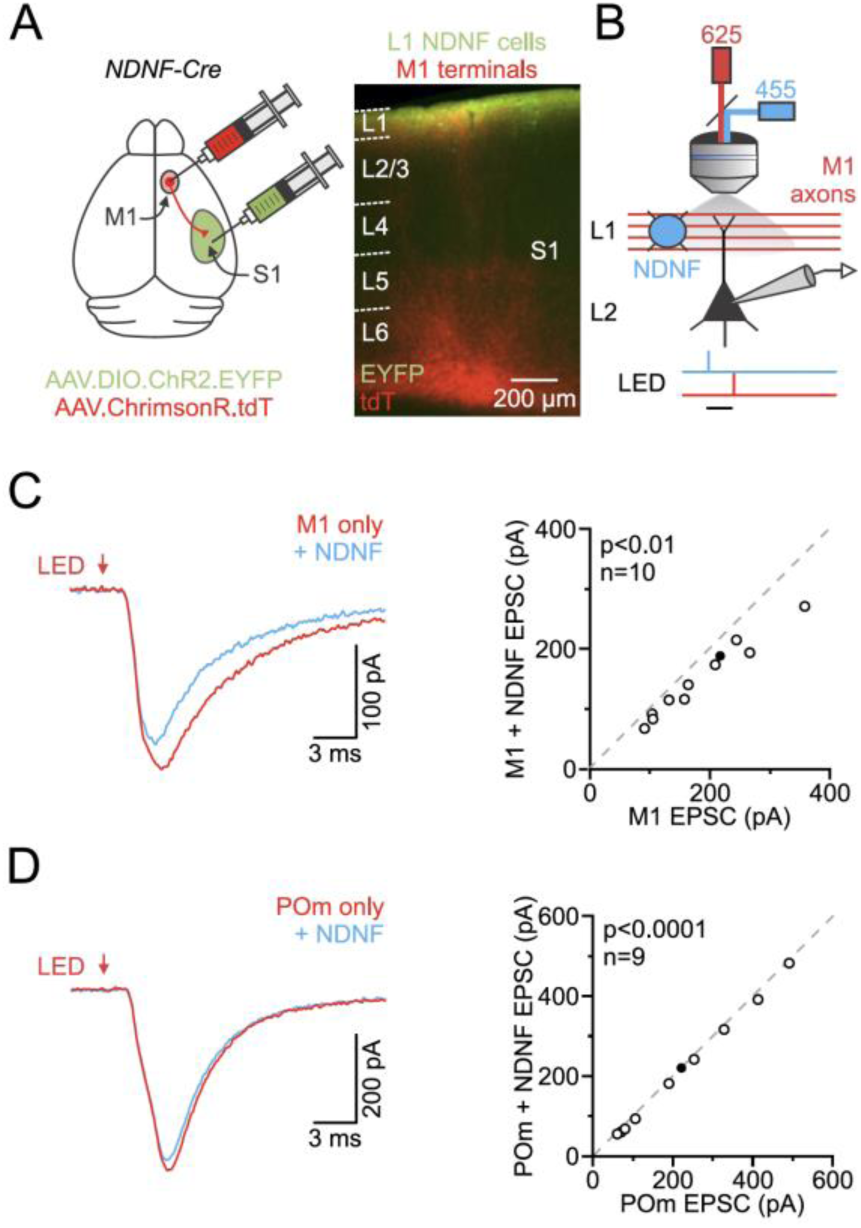
L1 NDNF cell activity inhibits optically-evoked top-down corticocortical and thalamocortical synapses via GABA_B_ receptor activation. **(a)** Left, schematic of an *NDNF-Cre* mouse brain showing a dual viral injection into the M1 and ipsilateral S1 of a Cre-independent AAV.ChrimsonR.tdT and a Cre-dependent AAV.DIO.ChR2.EYFP, respectively. Right, epifluorescent image of a fixed brain section at the level of S1 showing the overlap of ChR2/EYFP-expressing NDNF cells and ChrimsonR-tdT-expressing M1 axons/terminals in L1. **(b)** Schematic of the optical stimulation and recording configuration for dual-color optogenetic experiments in S1 of the *NDNF-Cre* mouse. Shown is the optical stimulation of L1 ChR2-expressing NDNF cells (blue, with 455 nm LED) and ChrimsonR-expressing M1 axons (red, with 525 nm LED), as well as whole-cell recording from a non-expressing L2 pyramidal cell. We recorded while alternating between control red light stimulation trials (0.5 ms, 625 nm) and trials in which we presented the blue light stimulation (0.5 ms, 2 mW, 455 nm) 50 ms before red light stimulation (0.5 ms, 625 nm). Horizontal scale bar, 50 ms. **(c)** Left, average M1 synaptic response in a L2 pyramidal cell of S1 evoked by a brief single red light stimulus in control conditions (red) and when NDNF cells were activated by a single blue light stimulus 50 ms before red light stimulation (blue). Right, summary graph showing the control synaptic response plotted against the response with NDNF cell activation (n = 10 cells, 5 mice). The filled black circle represents the mean. **(d)** Same as **(c)** but of the effects of blue light activation of NDNF cells on red-light evoked POm synaptic responses (n = 9 cells, 3 mice).

Since ChrimsonR is activated to some extent by blue light^91^, we first confirmed the suitability of this strategy by performing single opsin controls to determine the light intensities necessary to evoke reliable blue (455 nm) light-driven spiking in ChR2-expressing NDNF neurons without activating ChrimsonR-expressing axons/terminals. We observed clear differences in sensitivity to brief, low-intensity blue light stimulation (0.5 ms, 2 mW, 1.77 mW/mm^2^), with 68.8 ± 12.0% of ChR2-expressing NDNF cells responding with action potentials (n = 16 cells, 3 mice; **Fig. S2a**). In contrast, the same low-intensity blue light resulted in no evoked synaptic responses from ChrimsonR-expressing axons (n = 8 cells, 2 mice; **Fig. S2a**). Synaptic responses could be evoked from ChrimsonR-expressing axons/terminals by blue light stimulation, but this required intensities greater than or equal to 16 mW (9.02 mW/mm^2^). Furthermore, we did not observe ChR2-expressing NDNF cells respond to red light stimulation (**Fig. S2b**), confirming the suitability of our strategy.

To determine how NDNF cell activation affects M1 inputs, we recorded synaptic responses at L2 pyramidal neurons in the presence of the GABA_A_ receptor antagonist, picrotoxin (50 µM), and alternated control trials (0.5 ms, 625 nm red light only) with trials in which NDNF cells were briefly (0.5 ms, 2 mW) photostimulated with blue light 50 ms before red light stimulation of M1 axons/terminals (**Fig. 4b**). Using this approach, we found that NDNF cell activation significantly suppressed the amplitude of M1-evoked EPSCs (19.4 ± 2.0%; n = 10 cells, 5 mice; p = 0.00111, paired sample t-test; **Fig. 4c**). This suppressive effect was absent in animals not injected with ChR2 (n = 6 cells, 2 mice; p = 0.15, paired sample t-test; data not shown). Moreover, application of CGP-55845 (4 µM) prevented the NDNF-mediated suppression of M1 inputs, indicating the involvement of GABA_B_ receptor activation in the regulation of M1 synaptic strength (Control: 21.6 ± 4.3%, +CGP: 6.7 ± 0.9%, n = 3, 3 mice; **Fig. S2e**). Consistent with GABA_B_ receptor activity^60,63^, we also observed that NDNF-mediated regulation of M1 synapses was long-lasting, with significant inhibitory effects occurring with 200 ms intervals (**Fig. S2d**). Performing the same dual-color optogenetic experiments with ChrimsonR-expressing POm axons resulted in a much smaller modulation after NDNF cell activation (8.5 ± 1.8%, n = 9, 3 mice; p = 3.30628 x 10^-5^, paired sample t-test; **Fig. 4d**), which was not present in a control, non-ChR2 injected animal (n = 4, 1 mouse; p = 0.69494, paired sample t-test). The weaker NDNF-mediated modulation of POm synapses is consistent with the earlier observation that these synapses are less sensitive to baclofen and GABA_B_ receptor activation.

Overall, these data clearly show that GABA release from NDNF interneurons is sufficient to inhibit top-down synapses in L1 by activating presynaptic GABA_B_ receptors. Furthermore, the results demonstrate that their output attenuates top-down corticocortical input more than higher-order thalamic input, highlighting a critical role of L1 NDNF cells in modulating corticocortical communication.

### SOM cells contribute less to presynaptic modulation of L1 top-down synapses

SOM-expressing interneurons are also known to have ascending axons that arborize significantly in L1 of the neocortex^93–95^, and this class has previously been shown to engage GABA_B_ receptors^96,97^, suggesting that they may also modify transmission of top-down corticocortical information in L1. To test this idea, we injected AAV-ChrimsonR-tdT into M1 and AAV-DIO-ChR2-EYFP into S1 of *SOM-Cre* mice and repeated the previously described dual-color optogenetic experiment (**Fig. 5a and 5b**). In control experiments, we confirmed that low-intensity blue light (2 mW) reliably evoked spiking in ChR2-expressing SOM cells, and that both a single blue light stimulus and a 20 Hz optical train were sufficient to evoke GABA_B_-mediated responses (**Fig. S3a and S3b**). When we performed the dual-color optogenetic experiment, with picrotoxin (50 µM) in the bath, we found that SOM cell activation with a single stimulus resulted in a small but significant suppression of the amplitude of M1-evoked EPSCs (5.9 ± 1.4%, n = 8 cells, 4 mice; p = 0.01919, paired sample t-test; **Fig. 5c**). In contrast, activating SOM cells with a 20 Hz optical train (5 pulses) produced a stronger inhibition of M1-evoked synaptic responses (12.4 ± 2.0%, n = 8 cells, 4 mice; p = 2.86 x 10^-4^, paired sample t-test; **Fig. 5d**). However, even the SOM-mediated suppression generated by these optical trains was notably weaker than that produced by NDNF cells with a single stimulus (p = 0.02755, two-sample t-test). Altogether, these findings indicate that local SOM-expressing interneurons can also inhibit the synaptic strength of top-down corticocortical input, and that the magnitude of inhibition varies with frequency.

**Fig 5.**
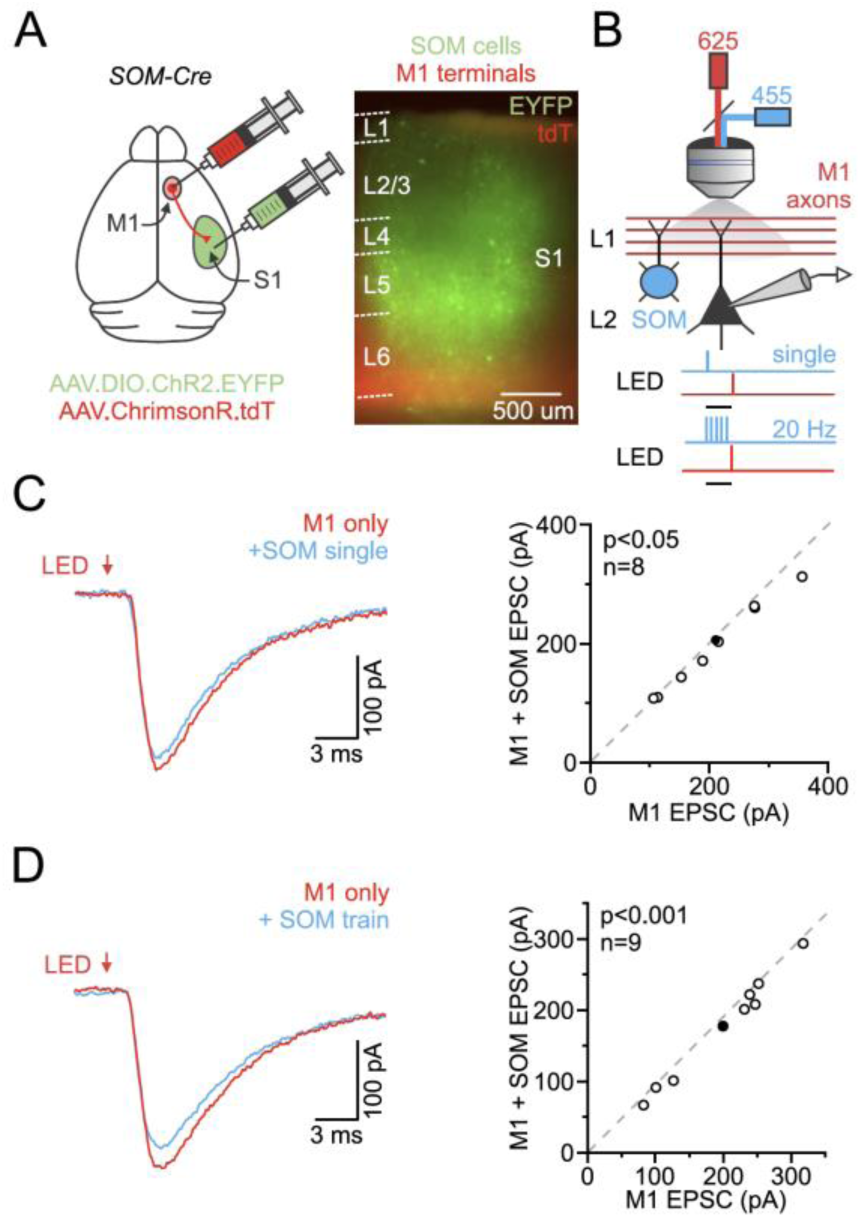
SOM cell activity is less capable of inhibiting the synaptic strength of M1 inputs, and the level of suppression varies depending on the frequency. **(a)** Left, schematic of a SOM-Cre mouse brain showing a dual injection into the M1 and ipsilateral S1 of a Cre-independent AAV.ChrimsonR.tdT and a Cre-dependent AAV.DIO.ChR2.EYFP, respectively. Right, epifluorescent image of a live brain slice at the level of S1 showing ChR2/EYFP-expressing SOM cells and ChrimsonR-expressing M1 axons/terminals. **(b)** Schematic of the optical stimulation and recording configuration for dual-color optogenetic experiments in the S1 of the *SOM-Cre* mouse. Shown is the optical stimulation of ChR2-expressing SOM cells (blue, with 455 nm LED) and ChrimsonR-expressing M1 axons (red, with 525 nm LED), as well as whole-cell recording from a non-expressing L2 pyramidal cell. We recorded while alternating between control red light stimulation trials (0.5 ms, 625 nm) and trials in which we presented the blue light stimulation (0.5 ms, 2 mW, 455 nm) 50 ms before red light stimulation (0.5 ms, 625 nm), either as a single pulse or as a 20 Hz train (5 pulses). Horizontal scale bars, 50 ms (top) and 250 ms (bottom). **(c)**, Left, average M1 synaptic response in a L2 pyramidal cell of S1 evoked by a brief single red light stimulus in control conditions (red trace) and when SOM cells were activated by a single blue light stimulus 50 ms before red light stimulation (blue). Right, summary graph showing the control synaptic response plotted against the response with SOM cell activation (n = 8 cells, 4 mice). The filled black circle represents the mean. **(d)** Same as **(c)** but of the effects of a 20 Hz blue light activation of SOM cells on red-light evoked M1 synaptic responses (n = 8 cells, 4 mice).

### POm activity drives L1 NGF cells

Our data indicate that L1 NDNF cell activity inhibits the synaptic strength of M1 inputs to S1 more strongly than POm inputs via presynaptic GABA_B_ receptor activation. While previous studies show that NDNF cells in other cortices receive long-range inputs from various cortical and thalamic regions^90,98–100^, it remains unclear whether M1 or POm inputs effectively drive NDNF cell spiking in S1, with important implications for how this L1 microcircuit is engaged. To better understand how M1 and POm engage this microcircuit, we next examined the effects of their inputs on molecularly distinct L1 NDNF cell subtypes.

Previous work has reported that L1 NDNF cells can be divided into two subtypes^62^. The first expresses neuropeptide Y (NPY), has wide, dense axonal arbors, and a late-spiking (LS) phenotype, consistent with NGF (neurogliaform) cells, which mediate long-lasting GABA_B_ inhibition following a single action potential^60,62,63^. The second subtype, called Canopy cells, lacks NPY, exhibits less-dense elongated axonal arbors, displays a non-late-spiking (NLS) phenotype, and is unable to produce a unitary GABA_B_ response. In contrast, other work in the auditory cortex reported no correspondence with NPY co-expression^18^. To determine whether NPY is a good marker for defining subpopulations of NDNF cells in our preparation, we crossed the *NDNF-Cre* line with the NPY-hrGFP mouse (**Fig. 6a**). We subsequently labeled NDNF-expressing cells by injecting S1 with a Cre-dependent mCherry virus. Whole-cell recordings from NDNF/NPY+ and NDNF/NPY-cells revealed distinct firing patterns consistent with those previously described for NGF and Canopy cells^62^ **Fig. 6b**). Most NDNF/NPY+ (85.7%; 18 of 21 cells) neurons had an LS firing pattern at threshld (rheobase) currents, whereas nearly all NDNF/NPY-(90.5%; 19 of 21 cells) neurons displayed non-LS patterns (**Fig. 6c**). Moreover, these two NDNF-expressing groups showed other distinct electrophysiological properties that correlate well with those previously described for NGF and Canopy cells (hereafter called), including sag amplitude, afterhyperpolarization amplitude, and input resistance (**Table S1**)^62^. These data indicate that NPY expression is a good marker for defining distinct subtypes of NDNF cells.

**Fig 6.**
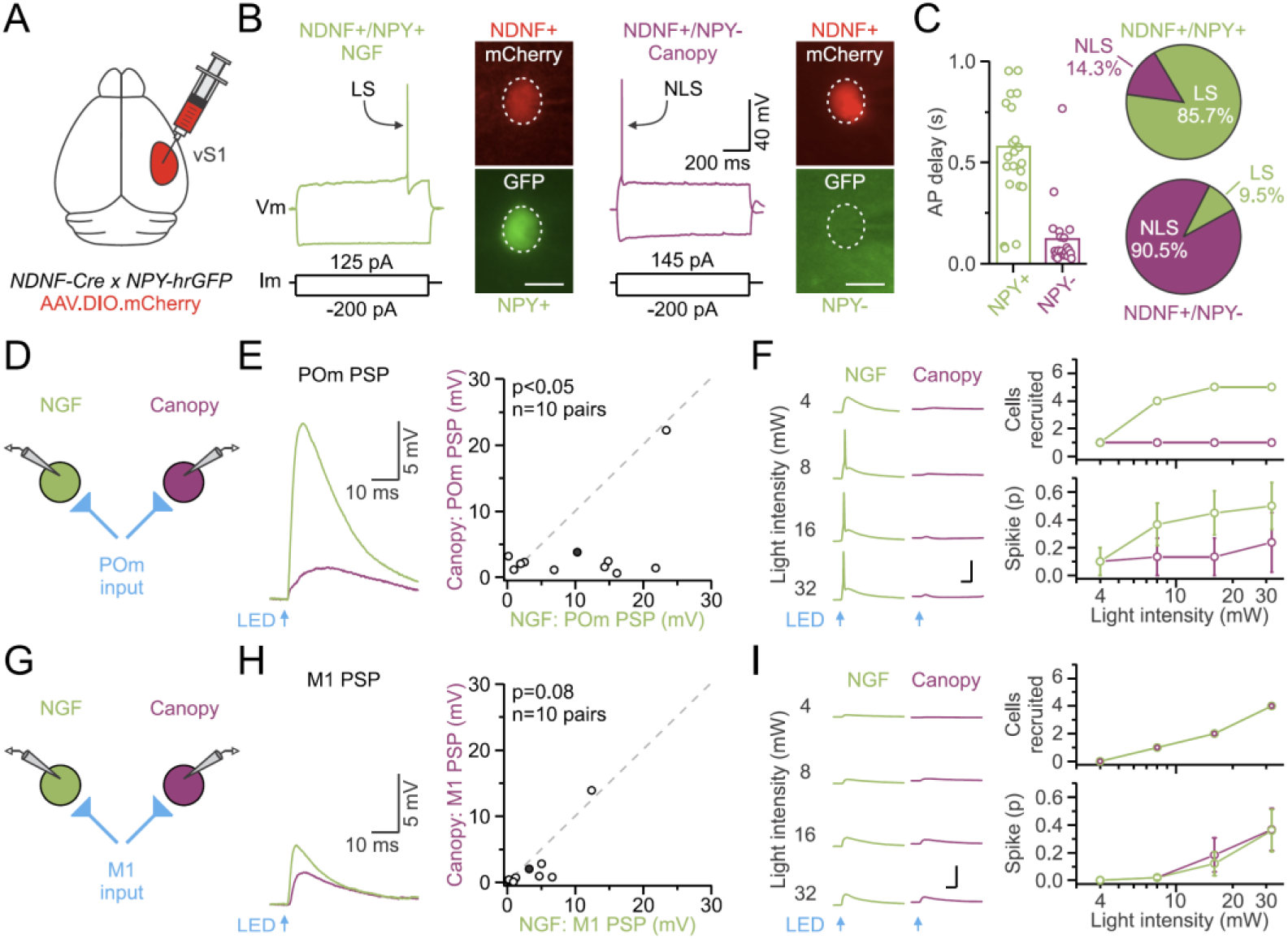
POm preferentially recruits L1 NGF cells. **(a)** Illustration of the genetic and viral approach used for targeting subtypes of L1 NDNF cells. **(b)** Voltage responses from an NDNF+/NPY+ (NGF: green) and an NDNF+/NPY-(Canopy: magenta) cell to injected current. The spiking response to a positive rheobase current was delayed in the NDNF+/NPY+ (NGF) cell, consistent with the late spiking (LS) phenotype. The NDNF+/NPY-(Canopy) had a non-late spiking (LS) phenotype in response to a rheobase current injection. Fluorescent images of the cells are also shown. Horizontal scale bars, 10 µm. **(c)** Population data comparing the mean delay to spike at threshold (rheobase) current (left), and pie charts displaying the proportions of NDNF/NPY+ and NDNF/NPY-cells with LS and non-LS firing patterns (right). **(d)** Schematic of the recording configuration for recording POm inputs onto pairs of L1 NGF (green) and Canopy (magenta) cells in S1. **(e)** Left, average M1-evoked PSP recording in an NGF-Canopy cell pair in current-clamp. Right, population data comparing the peak POM-evoked PSP in NGF cells versus Canopy cells (n = 10 pairs, 5 mice). The filled black circle represents the mean. **(f)** Left, example recordings from an NGF-Canopy cell pair evoked by POm stimulation of increasing intensities (4 - 32 mW). Right, summary graphs showing the number of NGF and Canopy cells recruited to spike (top) and spike probability (p; bottom) by stimulating POm axons/terminals at varying intensities. **(g)** Same as **(d)** but for recording M1 inputs. **(h)** Same as **(e)** but for M1-evoked PSPs recorded in NGF-Canopy cell pairs (n = 10 pairs, 5 mice). The filled black circle represents the mean. **(i)** Same as **(f)** but for NGF-Canopy cell pairs recorded during M1 stimulation (n = 10 pairs, 5 mice).

After validating our approach to access subtypes of L1 NDNF cells, we injected a Cre-independent AAV-ChR2-EYFP into POm or M1 of NDNF/NPY-hrGFP mice and performed sequential paired whole-cell recordings from neighboring L1 NGF and Canopy cells in the same slice (**Fig. 6d and 6g**). This strategy ensures localization within shared terminal arbors and controls for the variability in ChR2 expression across different slices and animals. In pairs studied this way, we found that low-intensity light stimulation (4 mW) of POm inputs elicited short-latency (∼2 ms) postsynaptic potentials (PSPs) in NGF cells that were larger than those in Canopy cells (NGF: 10.3 ± 2.8 mV; Canopy: 3.8 ± 2.1 mV; n = 10 pairs, 5 mice; p = 0.04883, Wilcoxon signed-rank pair test; **Fig. 6e**). In contrast, we found weaker, comparable M1-evoked PSP amplitudes across cell subtypes (NGF: 3.2 ± 1.3 mV; Canopy: 2.0 ±1.3 mV; n = 10 pairs, 5 mice; p = 0.08398, Wilcoxon signed-rank pair test; **Fig. 6h**). As we increased the light intensity, we observed that POm stimuli recruited NGF cells to spike at low light intensities (8 mW: 4/10 cells) but rarely elicited spiking in Canopy cells (8 mW: 1/10 cells) (**Fig. 6f**). At the same intensity, M1 input rarely recruited action potentials in either cell type (**Fig. 6i**). Overall, across pathways and cell subtypes, POm exerted a greater influence on the spiking activity of the NGF subtype across a wide range of light intensities. In contrast, M1 showed no significant preference or propensity to drive spiking of either subclass at low or moderate light intensities. Together, these results suggest that activity in the POm can activate presynaptic GABA_B_ receptors in L1 by driving spiking in L1 NGF cells.

## DISCUSSION

Here, we examined how GABA-mediated inhibition shapes top-down synaptic transmission in L1 of S1. Our results reveal that presynaptic GABA_B_ receptor activation suppresses transmitter release from top-down M1 and S2 cortical synapses more than from higher-order POm synapses. We also observed that the magnitude of GABA_B_ modulation varies by target cell type, being notably stronger at M1 synapses terminating on excitatory pyramidal cells than on GABAergic interneurons. Furthermore, we demonstrate that NDNF-expressing inhibitory cells effectively modulate top-down signaling, while SOM-expressing interneurons were less effective, with the magnitude of SOM-mediated suppression varying with firing frequency. Lastly, we identify L1 NGF cells as key targets of high-order thalamic projections, with POm stimulation driving these cells to spike. Collectively, our findings reveal a circuit in which the higher-order thalamus influences top-down corticocortical communication in S1 by recruiting L1 NGF cells, which, in turn, preferentially suppress glutamatergic transmission from top-down cortical synapses by activating presynaptic GABA_B_ receptors.

Previous studies have established the presence of GABA_B_ receptors in the cortex^101–110^, yet isolating their role in presynaptic modulation at long-range connections is challenging with conventional electrophysiological methods, primarily due to neighboring axons. In this study, we used AAVs to express the light-sensitive cation channel ChR2 in specific top-down cortical and thalamic projection neurons, enabling selective optical stimulation of their L1-targeting axons/terminals and investigation of how GABA_B_ receptor activation and specific cell types influence transmission. Similar optogenetic approaches have recently been used to study presynaptic modulation at other CNS synapses^54,61,111^, highlighting the potential of optical stimulation as a versatile tool for investigating presynaptic modulation at synapses that are not amenable to study using conventional methods. As L1 is also a key target of various subcortical neuromodulatory centers^17,19^, investigating how L1 top-down synapses are regulated by these neuromodulators warrants future studies, with significant implications for brain state control and cortical processing.

Our study clearly demonstrates that the activation of GABA_B_ receptors plays a critical role in modifying top-down signaling in S1. By blocking baclofen’s direct actions on the postsynaptic neuron with intracellular Cs^+^, we observed increases in both PPR and CV, strongly indicating that baclofen acts at presynaptic sites. At many glutamatergic synapses, GABA_B_ receptors containing the GABA_B1a_ subunit are preferentially targeted to presynaptic terminals^109,112–114^. When activated, these receptors are generally thought to inhibit voltage-gated Ca^2+^ channels, thereby reducing Ca^2+^ influx and the probability of synaptic release^49,51–53,115,116^. However, GABA_B_ receptors may also suppress release by shunting presynaptic action potentials through activation of K^+^ channels or by interacting with the vesicular release machinery^117,118^. Future studies will be important to determine the underlying mechanisms at these synapses.

L1 activity is influenced by two primary sources of glutamatergic afferent connections^61,81,99,100,119–121^: long-range corticocortical and thalamocortical, each conveying different information^17,19^. Our results demonstrate that these two distinct excitatory synapses are differentially affected by presynaptic GABA_B_ receptor activation. For example, 2.5 μM baclofen reduced EPSC amplitudes by 50% at corticocortical S2 synapses and 36% at corticocortical M1 synapses, while thalamocortical POm EPSCs only decreased by 18%. These findings not only indicate that top-down corticocortical afferent inputs are more sensitive to GABA_B_ receptor activation, but they also suggest a mechanism for adjusting the relative strength of excitatory top-down inputs during states when local GABA levels are high. The differential regulation of L1 synapse types also aligns with previous studies showing that local intracortical synapses are more sensitive to GABA_B_ modulation than sensory thalamocortical synapses^106,107^ and together imply that differential modulation of synapses is a key feature of the neocortex.

GABA_B_ receptor activation also influences the frequency sensitivity of these synapses. Long-range corticocortical responses typically show facilitation^81,122^, whereas POm responses exhibit depression. Consequently, activation of presynaptic GABA_B_ receptors may help minimize frequency-dependent depression of POm synapses, enabling more sustained transmission. Simultaneously, their activation may enhance the high-pass filtering imposed by facilitating synapses^123^, helping reduce low-frequency corticocortical communication while allowing higher-frequency signals to pass. Overall, our findings indicate that local inhibition plays a dynamic role in fine-tuning the strength and temporal response dynamics of distinct L1 afferents.

We found that GABA_B_-mediated suppression of M1 corticocortical synapses is also greater on pyramidal neurons than on inhibitory interneurons, indicating that GABA_B_ effects on excitatory M1 transmission are target cell-specific. Target-specific modulation of presynaptic release has been reported in various areas of the CNS^59,124–127^, and the mechanisms underlying this phenomenon include differential receptor expression in terminals within the same axon, mechanisms that suppress neurotransmitter release, and differences in local uptake mechanisms^124,128,129^. In future studies, it will be necessary to investigate these possibilities and to determine if other top-down pathways exhibit similar target cell-specific modulation. One potential physiological consequence of this biased modulation could be an increase in the responsiveness of inhibitory interneurons to top-down signaling relative to pyramidal neurons. However, due to cell-type-specific differences in long-range corticocortical circuitry, it might lead to different effects on the local circuit, depending on the specific pathway and the interneuron subtype recruited, and whether it results in disinhibition or feedforward inhibition^47,65,81,130,131^.

Our results highlight the crucial role of NDNF-expressing interneurons in the presynaptic modulation of top-down synapses in L1 of S1. These inhibitory cells, which belong to the 5HT3aR-expressing class of interneurons, are primarily found in cortical L1^62,90^ and have been reported to comprise at least two subtypes: NGF and Canopy cells, which differ in their NPY expression, morphology, and physiology^62,100^. Notably, NGF cells can elicit GABA_B_ responses in postsynaptic targets with a single action potential^62,63,132^, whereas Canopy cells do not^62^. This unique capability is linked to NGF cell axons, which have numerous release sites not associated with classic synaptic junctions, resulting in GABA volume transmission^60^. Consistent with previous reports^62^, our data indicate that most L1 NDNF-NPY-expressing neurons correspond to NGF cells based on their LS firing pattern. However, a recent study in the auditory cortex has suggested that NPY expression does not define distinct sub-classes^18^. Area differences could explain this discrepancy, but they may also reflect age-related changes in NPY, as previously suggested^18,133^.

Optogenetic activation of NDNF cells can inhibit the synaptic strength of top-down inputs for several hundred milliseconds, an effect blocked by CGP-55845, indicating activation of presynaptic GABA_B_ receptors. Note that the inclusion of picrotoxin in the experiments ruled out GABA_A_ receptors as contributing to presynaptic modulation^134^. Our findings also indicate that NDNF output is far less effective at suppressing higher-order POm afferents than M1, despite the close intermingling of top-down terminals within L1 (however, see^119,135^), which challenges the prevailing belief that NGF cells provide broad, non-selective inhibition via GABA volume transmission^60^. This aligns well with findings that L4 NGF cells selectively inhibit PV cell output while sparing nearby thalamocortical transmission^59^, further supporting the idea that some NGF cells have a more restricted spatial influence. This work also implies that local GABA release from NGF cells can alter the information encoded at individual L1 synapses, independent of what is conveyed at the soma and perhaps in collaterals of the same axon that target deeper layers within the same region (e.g., L5/6)^64,131,136^, or even different cortical areas^137^.

In contrast, optogenetic activation of SOM interneurons is far less effective at inhibiting L1 top-down synapses. SOM cells, a major class of cortical interneuron that target L1 tuft dendrites of pyramidal neurons^94,138–140^, have synapses that are initially less reliable but become more effective with repeated activation^141^. Consistent with this, we found that they suppress L1 inputs more effectively during repetitive stimulation, a condition necessary to activate GABA_B_ receptors^128,129^. Additionally, SOM cells are more sensitive to local inputs than long-range afferents^65,81,131,142,143^, and these excitatory connections display short-term facilitation^141^, making these inhibitory cells quite unresponsive to transient stimuli but highly responsive to sustained high-frequency activity within the local circuit^41,141,142,144^. NGF synapses, on the other hand, cannot maintain output during repetitive firing^63^. One possibility is that these two interneurons may act in mutually exclusive manners, modifying the transmission of top-down information across different cortical processing states that reflect ongoing behavioral demands. Indeed, it has been reported that the sensory-evoked responses, as well as the experience- and state-dependent aspects of NDNF and SOM responses, differ markedly^90,98^. However, SOM cells have also been shown to directly suppress the activity of NDNF cells, further complicating these circuits^90^.

Although L1 is a primary target for long-range corticocortical and higher-order thalamic projections, few studies have examined how top-down inputs engage specific cell types in L1 of S1^131^. Our findings indicate that corticocortical inputs from M1 engage NDNF cells. However, while both NGF and Canopy cells exhibit similar response strengths to M1 stimulation, they were rarely driven to fire action potentials. Thus, M1 does not seem to be a strong driver of L1 NGF cells. In contrast, responses to the POm were significantly stronger in NGF cells than in Canopy cells, and in half the cases, elicit action potentials in these cells at low light intensities. The observation that not all NGF cells responded with spikes to POm stimuli could be due to strong lateral inhibition between adjacent NDNF cells^145^ or variable ChR2 expression. Nevertheless, POm appears to be a strong driver of L1 NGF cells in S1, a finding that aligns with results from other studies of auditory and frontal cortices^61,99^.

Based on these findings, we propose that the higher-order thalamus plays a crucial role in suppressing top-down corticocortical signaling to L1 by recruiting NGF cells. These cells activate presynaptic GABA_B_ receptors, which decrease the synaptic strength of corticocortical input. This presynaptic circuit aligns with emerging insights regarding the roles of NDNF/NGF cells in tightly regulating pyramidal cell excitability. They do this by directly inhibiting the distal apical tuft dendrites of L2/3 and L5 neurons through both GABA_A_ and GABA_B_ receptors^62,90,146–148^, thereby controlling complex dendritic spikes in these neurons^99,110,148^. Additionally, they appear to indirectly disinhibit the somata of these cells by inhibiting PV-expressing interneurons^18,99,149^. Taken together, these studies and our study suggest that the higher-order thalamus may shift the information-processing strategy of pyramidal cells from integrating top-down and bottom-up signals to one that enhances bottom-up sensory responsiveness.

Importantly, previous studies have also demonstrated that POm neurons directly target VIP interneurons^131,150,151^, which in turn disinhibit the distal apical tuft dendrites of pyramidal neurons by inhibiting dendritic-targeting SOM cells^45^. This raises the intriguing possibility that the POm engages two distinct inhibitory microcircuits that can influence the dendritic and somatic activity of S1 excitatory neurons, with significant implications for somatosensory processing. Supporting this hypothesis, Anastasiades et al. (2021) showed in the prefrontal cortex (PFC) that two distinct thalamic nuclei differentially affect PFC excitability through layer 1 NDNF and VIP microcircuits. In the future, it will be important to determine whether the engagement of these two inhibitory circuits is mediated by distinct POm subpopulations that compete under different contexts to influence top-down signaling^78,152,153^, whether they influence different cortical excitatory cells^154,155^, or whether different patterns of POm activity recruit each inhibitory circuit^63,131^.

The POm is a higher-order thalamic structure that is reciprocally connected with multiple somatosensory and motor cortical areas and receives input from many other subcortical regions^156–160^, and thus is well positioned to influence cortical activity. Despite its strategic position and widespread innervation, our understanding of the functions of POm and other higher-order nuclei still lags behind that of the core sensory nuclei^161^. While research supports roles in behavioral flexibility, such as learning, perception, and behavioral salience, recent *in vivo* studies suggest that POm and other higher-order nuclei convey non-sensory, contextual signals about internal state to the cortex^35,36,69,162^. Supporting this idea, NDNF cells show heightened activity and sensory responsiveness during arousal^98,163^. Therefore, during arousal, POm may provide L1 NGF cells with contextual information about the ongoing internal state. This communication could help reconfigure cortical circuits to suppress top-down corticocortical signals and selectively enhance bottom-up sensory-evoked activity^164^ in accordance with the global brain state.

In a seminal study by Pardi et al. (2020), it was demonstrated that NDNF interneurons inhibit L1 high-order auditory thalamic terminals involved in learning. Our findings help clarify that top-down corticocortical synapses are more sensitive to presynaptic GABA_B_ modulation than thalamic synapses, primarily decreasing transmitter release at terminals contacting pyramidal cells. Similar to Pardi et al. (2020), we also confirm that NDNF cells exert presynaptic control over top-down inputs in S1. In addition, we find that the higher-order thalamus drives NGF cells in L1. Together, our studies indicate that presynaptic modulation of L1 top-down afferents by NGF cells may be a conserved circuit motif across cortical regions with shared functions. The present study provides a better understanding of this circuit, aiding future studies investigating its potential role in neocortical processing.

## METHODS

### Animals

All procedures were carried out in accordance with the National Institutes of Health (NIH) Guidelines for the Care and Use of Laboratory Animals and approved by the Michigan State University Institutional Animal Care and Use Committee (IACUC). We used the following mouse lines in this study: Crl: CD1 (ICR) (Charles River: 022), Ai14 (Jackson Labs: 007908)^165^, PV-Cre (Jackson Labs: 008069)^88^, SOM-IRES-Cre (Jackson Labs: 013044)^89^, 5HT3a-EGFP (MMRRC: 000273-UNC)^166^, NPY-hrGFP (Jackson Labs: 006417)^167^, G42 (Jackson Labs: 007677)^168^, NDNF-Cre (Jackson Labs: 030757)^62^. Experimental PV and SOM mice were generated by crossing homozygous Cre mice with homozygous Ai14 reporter mice, yielding heterozygous mice for the indicated genes. Animals were group-housed with same-sex littermates in a dedicated animal care facility maintained on a 12:12 h light-dark cycle. Food and water were available *ad libitum*. We used both male and female mice in this study.

### Stereotactic Virus Injections

For many experiments, we used an adeno-associated virus (AAV2) that encoded genes for hChR2 (H134R)-EYFP fusion proteins (rAAV2/hSyn-hChR2[H134R]-eYFP-WPREpA, AV4384, University of North Carolina (UNC) Viral Vector Core; titer = 3.5 x 10^12^ viral genomes(vg)/mL). To fluorescently label L1 NDNF cells, we used rAAV1/Ef1a-DIO-mCherry (Addgene; titer = 2.0 x 10^12^ vg/mL). For dual opsin experiments, we used an AAV2 virus that encoded genes for ChrimsonR-tdT fusion proteins (rAAV2/Syn-ChrimsonR-tdT, AV6096, UNC Viral Vector Core; titer = 3.7 x 10^12^ vg/mL) and ChR2-EYFP fusion proteins (rAAV1/Ef1a-DIO-hChR2(H134R)-eYFP, V17648, Addgene; titer = 2.0 x 10^12^ vg/mL).

For each surgery, the virus was injected unilaterally into M1, POm, S2, or S1 of mice *in vivo*, as previously described^81,136,169^. Injections were typically performed on mice ∼5 weeks of age (Mean injection age: 35 ± 1.3 days, range 18-64 days). Briefly, mice were anesthetized using a Ketamine-Dexmedetomidine cocktail diluted in sterile saline (KetaVed, 70-100 mg/kg; Dexdomitor, 0.25 mg/kg; intraperitoneally). Once deeply anesthetized, mice were secured in a digital stereotaxic frame with an integrated warming base to maintain core body temperature (Stoelting). Ophthalmic ointment was applied to the eyes to prevent drying during surgery (Patterson Veterinary Artificial Tears). Next, fine surgical scissors were used to make an incision over the skull, the scalp and periosteum were retracted, and a small craniotomy was made over the target site. A small amount of virus was then pressure-ejected via a glass micropipette attached to a Picospritzer pressure system (0.1-0.2 µL per injection site over 5-10 min). After the injection, the pipette was held in place for an additional 5-10 min before being slowly advanced or withdrawn from the brain. The scalp was then closed with a surgical adhesive (GLUture) or sutures (Surgical Specialties, 1285B), and animals were given Buprenorphine-SR (0.05–0.1 mg/kg) subcutaneously for postoperative pain relief and antisedan (2.5 mg/kg) to reverse the effects of Dexdomitor. Mice recovered on a heating pad for a minimum of 1 h before returning to their home cage. *In vitro* experiments were typically conducted 2-3 weeks after viral injection to allow sufficient expression (Mean expression time: 20 ± 0.31, range 13-25 days). Coordinates from bregma for M1 were 1.25 mm lateral, 0.9 and 1.3 mm anterior, 0.40 and 1.0 mm depth. Coordinates from bregma for POm were 1.33 mm lateral, 1.35 mm posterior, and 2.93 mm depth. Coordinates from bregma for S2 were 4.2 mm lateral, 1.1 mm posterior, and 1.16 mm depth. Coordinates from bregma for S1 were 3.10 mm lateral, 0.9 mm posterior, and 0.35 mm depth.

### *In Vitro* Slice Preparation

Acute coronal brain slices (300 μm thick) containing S1 were prepared for *in vitro* recording as described previously^81,122,169^. Briefly, animals were deeply anesthetized using isoflurane before being decapitated. Brains were extracted and placed in a cold (∼4°C) oxygenated (95% O_2_, 5% CO_2_) slicing solution containing (in mM) 3 KCl, 1.25 NaH_2_PO_4_, 10 MgSO_4_, 0.5 CaCl_2_, 26 NaHCO_3_, 10 glucose, and 234 sucrose. Brain slices were cut using a vibratome (Leica, VT1200S) and then placed in a holding chamber with warm (32°C) oxygenated artificial cerebrospinal fluid (ACSF) containing (in mM): 126 NaCl, 3 KCl, 1.25 NaH_2_PO_4_, 2 MgSO_4_, 2 CaCl_2_, 26 NaHCO_3_, and 10 glucose. Slices were kept at 32°C for 20 min, then at room temperature for at least 40 min before recording. Slices containing the injection site were always collected and imaged to confirm the accuracy of the injection site and ensure tissue health. We excluded any mice with signs of tissue damage or off-target injections.

### *In Vitro* Electrophysiological Recordings and Data Acquisition

Individual brain slices were transferred to a submersion-type recording chamber and bathed continually (3 mL/min) with warm (32 ± 1°C) oxygenated ACSF containing (in mM): 126 NaCl, 3 KCl, 1.25 NaH_2_PO_4_, 1 MgSO_4_, 1.2 CaCl_2_, 26 NaHCO_3_, and 10 glucose. Neurons were visualized using infrared differential interference contrast optics (IR-DIC) with a Zeiss Axio Examiner.A1 microscope equipped with a 40x water immersion objective (Zeiss, W Plan-Apo 40x/1.0 NA) and video camera (Olympus, XM10-IR). For voltage-clamp experiments, whole-cell recordings were made using patch pipettes with tip resistances of 4-6 Megaohms when filled with a cesium-based internal solution containing (in mM): 130 Cs-gluconate, 4 CsCl, 2 NaCl, 10 HEPES, 0.2 EGTA, 4 ATP-Mg, 0.3 GTP-tris,14 phosphocreatine-di tris, 2 mM QX-314 (Tocris, Cat# 2313) and 15 mM Tetraethylammonium chloride (TEA; Sigma, Cat# T2265) (pH 7.25, 290 mOsm). For current clamp experiments, a potassium-based internal solution was used, containing (in mM): 130 K-gluconate, 4 KCl, 2 NaCl, 10 HEPES, 0.2 EGTA, 4 ATP-Mg, 0.3 GTP-Tris, and 14 phosphocreatine di-tris (pH 7.25, 290 mOsm). Voltages were corrected for a -14 mV liquid junction potential.

Electrophysiological data were acquired and digitized at 20 kHz using Molecular Devices hardware and software (MultiClamp 700B amplifier, Digidata 1550B4, pClamp 11). Before digitization, signals were low-pass filtered at 3 kHz (for voltage-clamp) or 10 kHz (for current-clamp). During recordings, the pipette capacitances were neutralized, and series resistances (typically 7-25 megaohms) were compensated online (70% compensation for voltage-clamp, 100% for current-clamp). Series resistances were continually monitored throughout recordings and adjusted during experiments to ensure sufficient compensation. Data was discarded if the series resistance changed more than 20% from the original value. Pharmacological agents included R-Baclofen (Tocris, Cat# 0796), CGP55845 hydrochloride (Tocris, Cat# 1248), and Picrotoxin (Sigma, Cat# P1675). All receptor antagonists were bath applied, whereas the agonist R-Baclofen was diluted in HEPES-buffered ACSF containing (in mM): 126 NaCl, 3 KCl, 1.25 NaH_2_PO_4_, 1 MgSO_4_, 1.2 CaCl_2_,10 glucose, 5 HEPES (pH 7.25, 300 mOsm) and administered via a syringe pump (KD-scientific).

### Photostimulation

ChR2 was stimulated using a high-power white light-emitting diode (LED) (Mightex, LCS-5500-03-22) driven by an LED controller (Mightex, BLS-1000-2). For the dual-opsin photostimulation experiments, ChR2 and ChrimsonR were excited using a high-power blue (455 nm; LCS-0455-03-22) and red (625 nm; LCS-0625-03-22) LED, respectively, combined by a multiwavelength beam combiner (Mightex, LCS-BC25-0495). LED on/off times were fast (<50 us) and of constant amplitude and duration when verified with a fast photodiode (Thorlabs, DET36A). The light was collimated and reflected through a single-edge dichroic beam-splitter (Semrock, FF660-FDi02) and a high-magnification water immersion objective (Zeiss, W Plan-Apo 40x/1.0 NA), resulting in an estimated spot diameter of ∼1500 um and maximum LED power at the focal plane of 45.10 mW (white), 57.0 mW (blue), 41.6 mW (red). Stimuli were delivered as 0.5-ms flashes every 10-30 seconds and were directed at ChR2 or ChrimsonR-expressing axons/terminals by centering the light over the recorded cell. For each pathway, the LED intensity was adjusted to evoke an EPSC of 100-300 pA in the recorded neuron when held at -75 mV in voltage-clamp. Optical stimulation was administered at 15-second intervals, with a stable 3-minute baseline period preceding each baclofen test. Baclofen was delivered via a syringe pump for 30 seconds, and responses were recorded for another 10 minutes.

### Live Slice Imaging

Before recording, all live brain slices were imaged using a Zeiss Axio Examiner.A1 microscope equipped with a 2.5x objective (Zeiss, EC Plan-Neofluar) and an Olympus XM10IR camera, and cellSens software. Brightfield and fluorescence images were obtained to confirm the accuracy of the injection, tissue health at the injection site, and overall expression in axons/terminals in S1. Live slices were maintained in a submersion recording chamber during imaging and continuously bathed with room-temperature oxygenated ACSF.

## QUANTIFICATION AND STATISTICAL ANALYSIS

### Electrophysiology

Analysis of electrophysiological data was performed using Molecular Devices Clampfit 11 and Microsoft Excel. Synaptic responses to optical stimulation were measured from postsynaptic neurons recorded in whole-cell voltage-clamp or current-clamp. The amplitude of an evoked response was measured relative to a baseline before the stimulus (10 ms). The peak amplitude was measured over 10 ms immediately following stimulus onset. The baseline period was defined as 3 minutes (12 trials) of stable baseline recording before the onset of baclofen. Peak modulation was calculated by averaging 10 sweeps following baclofen application. Paired-pulse ratios were measured using pairs of optical stimuli at 20 Hz and calculated by dividing the amplitude of the second pulse by the amplitude of the first pulse. The coefficient of variation (CV) was calculated as the standard deviation divided by the mean amplitude modulation. Most L2 and L5 pyramidal neurons were identified by their morphology under IR-DIC and subsequently by electrophysiology. All inhibitory cells were identified using specific transgenic mouse lines.

Intrinsic properties were analyzed as previously described^136^. The delay to spike was measured as the time from the start of the current injection (1 second) to the first AP at threshold (rheobase) current. Cells were classified as late spiking (LS) if delays were longer than 350 ms, and non-LS if delays were less than 350 ms. Sag percent was calculated by first determining the sag amplitude. This was done by finding the negative-going peak within the first 200 ms of the negative current injection (−200 pA, 1 second) and subtracting the average steady-state voltage measured during the last 200 ms of the current injection. After finding the sag amplitude, it was divided by the total voltage deflection during the first 200 ms to represent the percent sag relative to the total voltage.

### Statistical Analysis

No formal randomization method was used in this study, but all mice and cells recorded were selected at random. Experimenters were not blind to the experimental groups. The Shapiro-Wilk test was used to determine normality, followed by appropriate parametric (paired sample t-test or two-sample t-test) or nonparametric tests (Wilcoxon paired signed-rank test or Mann-Whitney U test), as indicated in the results. Additionally, a Kruskal-Wallis ANOVA was used to compare multiple groups with nonparametric distributions. All tests were also two-tailed. Statistical significance was defined as p<0.05, and data are represented as mean ± SEM.

## ACKNOWLEDGMENTS

This research was supported by NIH grants R01-NS117636 (to S.R.C).

## AUTHOR CONTIBUTIONS

S.R.C. and K.E.B. designed the experiments, K.E.B., M.L.R., G.R.G., L.X., and M.K.S. performed the viral injections, K.E.B. and M.L.R. performed the electrophysiological experiments, K.E.B. analyzed the data, S.R.C. and K.E.B. prepared and wrote the manuscript. All authors reviewed and approved the final manuscript.

## DECLARATION OF GENERATIVE AI AND AI-ASSISTED TECHNOLOGIES

None

## SUPPLEMENTAL FIGURES, TABLES, AND FIGURE LEGENDS

**Supplemental Fig 1, related to Fig 1.**
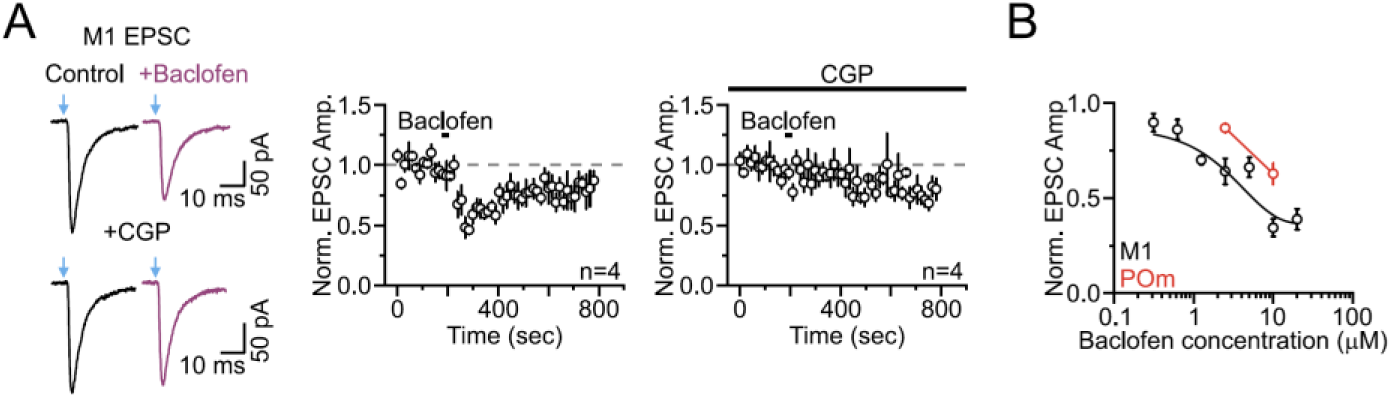
The effect of baclofen and CGP-55845 on M1 synaptic responses and the dose-response relationship of baclofen suppression on M1 and POm responses. **(a)** Left, average M1 synaptic response in a L2 pyramidal cell of S1 evoked by a single optical stimulus in control conditions (black) and after adding baclofen (2.5 µM, magenta), with and without CGP-55845 (4 µM) in the bath. Right, population data showing the average time course of baclofen effects (30 sec application) on M1 evoked EPSC amplitudes with or without CGP55845 in the bath (n = 4 cells, 3 mice). **(b)** Dose-response relationship of baclofen suppression on M1 (black) EPSC amplitudes: 0.31 µM (n = 5 cells, 2 mice), 0.63 µM (n = 5 cells, 2 mice), 1.25 µM (n = 4 cells, 2 mice), 2.5 µM (n = 13 cells, 9 mice), 5 µM (n = 7 cells, 4 mice), 10 µM (n = 4 cells, 4 mice), 20 µM (n = 6 cells, 4 mice). The effect of baclofen on POm (red) synaptic responses is also plotted for 2.5 µM (n = 8 cells, 3 mice) and 10 µM (n = 4 cells, 2 mice).

**Supplemental Fig 2, related to Fig 4.**
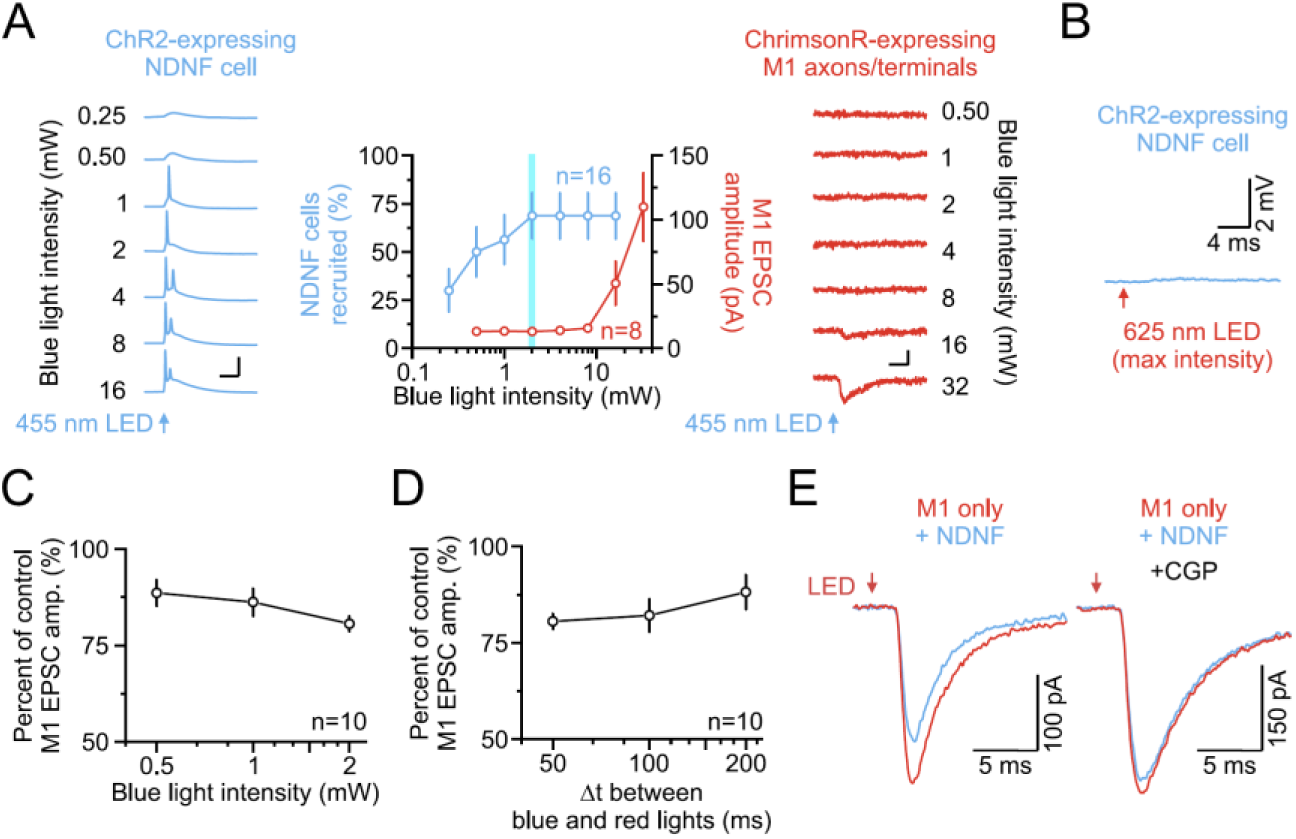
Independent optical excitation of ChR2-expressing interneurons and ChrimsonR-expressing L1 axons. **(a)** Left, example recordings from a single ChR2-expressing NDNF cell of S1 evoked by brief (0.5 ms) blue light stimulation of increasing intensities (0.25 - 16 mW shown. Right, example synaptic responses in a non-expressing L2 pyramidal cell of S1 evoked by brief (0.5 ms) blue light stimulation of increasing intensity over ChrimsonR-expressing M1 axons/terminals (0.5 - 32mW shown). Vertical scale bars, 40 mV (left) and 40 pA (right). Middle, summary plot showing the percentage of NDNF cells recruited and the amplitude of the M1 EPSC evoked by blue light stimulation (NDNF cells: n = 16 cells, 3 mice; M1 EPSCs: n = 8 cells, 3 mice). Note that at a 2 mW blue light intensity (blue vertical line), nearly 70% of ChR2-expressing NDNF cells responded, while there was no detectable synaptic response from ChrimsonR-expressing M1 axons. **(b)** ChR2-expressing NDNF cells did not respond to red light stimulation. Example response from a ChR2-expressing NDNF cell of S1 evoked by max red light stimulation (top: 42 mW, 0.5 ms). **(c)** Plot showing the effect of NDNF cell activation as the percentage of control M1 EPSC amplitude against the intensity of the blue light stimulation of ChR2-expressing NDNF cells. Control M1 EPSCs evoked by red light alone (n = 10 cells, 5 mice). **(d)** Plot showing the effect of NDNF cell activation as the percentage of control M1 EPSC amplitude against the interval (Δt) between the onset of the blue light and subsequent red light. Control M1 EPSCs evoked by red light alone (n = 10 cells, 5 mice). **(e)** Example recordings showing presynaptic inhibition of a M1 synaptic response with NDNF cell activation and then subsequent blockade of this suppression with a GABA_B_ receptor antagonist (4 µM; CGP-55845) in the bath (n = 3, 3 mice).

**Supplemental Fig 3, related to Fig 5.**
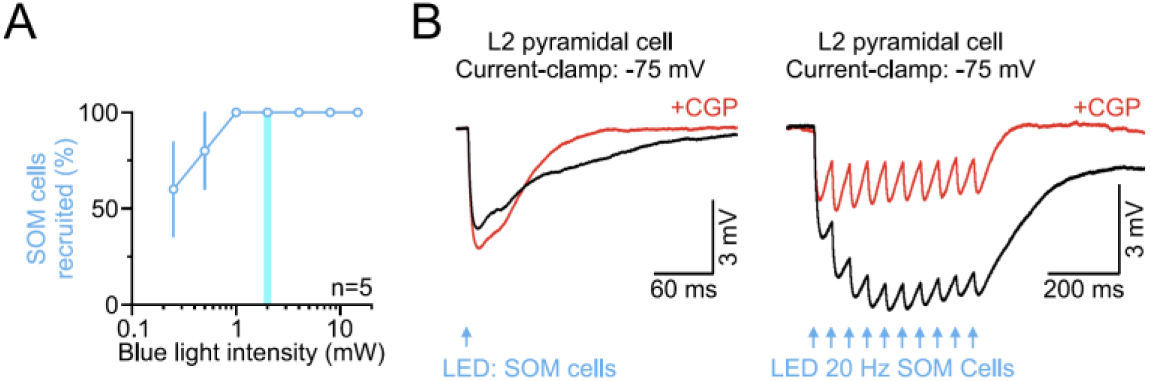
Low-intensity blue light stimulation reliably triggers action potentials in ChR2-expressing SOM cells and induces postsynaptic GABA_B_ responses. **(a)** Summary plot showing the percentage of SOM cells recruited to spike by blight light stimulation of increasing intensity from 0.25 to 16 mW (n = 5 cells, 4 mice). Note that at a 2 mW blue light intensity (blue vertical line), 100% of ChR2-expressing SOM cells responded. **(b)** Effects of a GABA_B_ receptor antagonist on light-evoked SOM responses of a non-expressing L2 pyramidal cell. In control conditions (black), when held at -75 mV in current-clamp, a single blue light stimulus (top) and 20 Hz optical train (bottom) evoked large inhibitory postsynaptic potentials (IPSPs) with a fast and slow phase. Bath application of the GABA_B_ receptor antagonist CGP-55845 (2 µM, red) blocked the slow phase of the IPSP.

**Supplemental Table 1.** Electrophysiological properties of L1 NDNF+/NPY+ (NGF) and NDNF+/NPY- (Canopy) cells in S1. Related to Figure 6

|  | L1 NDNF+/NPY+ (NGF) |  |  | L1 NDNF+/NPY- (Canopy) |  |  | NPY+ vs NPY- |  |
| --- | --- | --- | --- | --- | --- | --- | --- | --- |
|  | Mean | ± s.e.m. | n cells / mice | Mean | ± s.e.m. | n cells / mice | P | Test |
| Resting membrane potential (mV) | -83.30 | ± 0.62 | 21/11 | -80.29 | ± 0.79 | 21/11 | 0.004 | t test |
| Membrane resistance (MΩ) | 255.15 | ± 17.29 | 16/10 | 236.23 | ± 41.86 | 15/9 | 0.042 | MW U test |
| Membrane time constant (ms) | 11.80 | ± 0.76 | 16/10 | 11.35 | ± 0.89 | 15/9 | 0.465 | MW U test |
| Membrane capacitance (pF) | 46.80 | ± 1.80 | 16/10 | 58.55 | ± 6.11 | 15/9 | 0.068 | t test |
| Rheobase (pA) | 66.30 | ± 6.02 | 21/11 | 91.66 | ± 9.19 | 21/11 | 0.026 | t test |
| Spike threshold (mV) | -56.37 | ± 0.52 | 21/11 | -56.38 | ± 0.62 | 21/11 | 0.980 | MW U test |
| Delay to spike (ms) | 539.92 | ± 56.88 | 21/11 | 122.45 | ± 36.25 | 21/11 | 2.88 x 10<sup>-6</sup> | MW U test |
| Spike amplitude (mV) | 66.89 | ± 1.13 | 21/11 | 71.70 | ± 1.59 | 21/11 | 0.018 | t test |
| Spike half-width (ms) | 0.76 | ± 0.03 | 21/11 | 0.68 | ± 0.03 | 21/11 | 0.025 | MW U test |
| Afterhyperpolarization amplitude (mV) | 16.51 | ± 0.48 | 21/11 | 13.60 | ± 0.86 | 21/11 | 0.005 | t test |
| Max rate of rise (mV*ms) | 339.27 | ± 12.33 | 21/11 | 368.13 | ± 13.32 | 21/11 | 0.120 | t test |
| Max rate of decay (mV*ms) | -76.32 | ± 3.76 | 21/11 | -97.80 | ± 5.74 | 21/11 | 0.003 | t test |
| Sag % | 7.05 | ± 0.01 | 21/11 | 14.35 | ± 0.02 | 21/11 | 0.005 | MW U test |
| Depolarizing hump (mV) | 2.17 | ± 0.27 | 21/11 | 5.09 | ± 0.50 | 21/11 | 1.31 x 10<sup>-4</sup> | MW U test |

\* See methods for an explanation of how the electrophysiological parameters were defined/measured. Data shown as mean ± s.e.m.
\* All membrane potentials were corrected for a -14 mV liquid junction potential.

